# Paracoccin impacts on *Paracoccidioides brasiliensis* virulence and susceptibility to antifungal drugs by modifying the expression of genes related to remodeling of the yeasts cell wall

**DOI:** 10.1101/2022.07.05.498818

**Authors:** Nayla de Souza Pitangui, Fabrício Freitas Fernandes, Thiago Aparecido da Silva, Relber Aguiar Gonçales, Maria Cristina Roque-Barreira

## Abstract

The study of the paracoccin lectin (PCN) has provided knowledge about its role in the biology of *Paracoccidioides brasiliensis* and in the pathogenesis of paracoccidioidomycosis (PCM). In this context, PCN has proved to be a promising immunomodulatory agent for the exploration of vaccine target molecules and/or for diagnostic or therapeutic purposes. Previous investigations allowed establishing PCN as a factor of fungal virulence. However, the effect PCN exerts on the yeast’s resistance to antifungal pharmacological agents used to treat human PCM are not known. Therefore, this work characterizes the role of PCN functional duality on virulence and susceptibility of *P. brasiliensis* to antifungal drugs. We show that the PCN overexpression increases the virulence of *P. brasiliensis* yeasts in an alternative model of infection, induces high susceptibility *in vitro* and *in vivo* of *P. brasiliensis* yeasts to antifungal therapy, and impact reducing relative mRNA expression of genes encoding proteins related to cell wall degradation. Conversely, PCN silencing minimized the yeasts’ virulence in *Galleria mellonella*, correlates with the lowest susceptibility to treatment with antifungal agent *in vivo* and impact differently from the PCN overexpression on the relative expression of markers related to *P. brasiliensis* yeasts cell wall remodelling. Our study demonstrates the impact of endogenous PCN on the *P. brasiliensis* yeasts’ virulence *vs*. susceptibility to antifungal drugs, the fungal biology, and the relationship of the yeasts-host cells.

## INTRODUCTION

Paracoccin (PCN) is a contributory virulence factor for establishing *Paracoccidioides brasiliensis* infection. PCN was identified in the last decade as a bifunctional protein, whose lectin domain binds to N-acetylglucosamine and chitin [1,2], while the enzymatic domain exerts an N-acetylglucosaminidase (NAGase) activity. Localized on the surface of *P. brasiliensis* yeast cells, mainly in the budding regions, PCN plays a prominent role in fungal growth. Its enzymatic activity is associated with remodeling the fungal cell wall, thanks to its ability to hydrolyze multimeric structures (chitin) in monosaccharides (GlcNAc).

Regarding the importance of PCN for the pathogenesis of *Paracoccidioides spp*., some investigations revealed the immunomodulatory effect of this protein. This property was deeply investigated because of its potential application in designing alternative paracoccidioidomycosis (PCM) treatment or prevention strategies. It is currently known that the administration of native or recombinant PCN in mice confers protective immunity against *P. brasiliensis* experimental infection, established through (*i*) direct PCN interaction with N-glycans of TLR receptors; (*ii*) induction of macrophage M1 polarization, mediated by the interaction of PCN with TLR4 and; (*iii*) stimulation of immune response mediated by T helper 1 cells (Th1) [3-6].

Recent studies have analyzed the impact of silencing and overexpression of the PCN gene (PADG_03347) on the virulence of *Paracoccidioides* spp. Briefly, the data indicate that, relative to wild type (WT, wild type) strain, PCN silencing has relevant consequences. The in vitro detected effects are: (a) reduced frequency of individual yeast cells; (b) blocking the yeast-mycelium transition; and (c) augmented susceptibility of yeasts to the fungicidal activity of macrophages. In vivo effects were verified by infecting mice with PCN silenced yeasts; the consequent disease was characterized by a) reduced pulmonary fungal burden, b) reduced tissue damage, and c) reduced mortality rate [7]. On the other hand, PCN overexpression in *P. brasiliensis* yeasts, compared to WT yeasts, leads to in vitro consequences: a) a more efficient yeast-mycelium transition occurred in a reduced time lapsed, and b) augmented yeasts resistance to the fungicidal activity of macrophages. In vivo infection with transformants that overexpress PCN resulted in: a) increased pulmonary fungal burden; b) severe tissue injury; and c) increased mortality rate of infected BALB/c mice [8]. So, there is enough evidence that PCN plays a relevant role in the pathogenesis of PCM, allowing us to consider that PCN is a virulence factor for fungi of the *Paracoccidioides* genus.

The PCN impact on the virulence of yeasts of the *Paracoccidioides* complex is well established. However, we still do not know the effect PCN exerts on the yeast’s resistance to antifungal pharmacological agents used to treat human PCM.

## METHODS

### *P. brasiliensis* strains

*P. brasiliensis* wild-type strain (wt-PCN), PCN-overregulated strain (ov-PCN), PCN-downregulated strain (kd-PCN) and empty vector (EV) were maintained in brain heart infusion (BHI) solid or liquid media (Kasvi, São José dos Pinhais, Brazil) supplemented with 1% glucose at 37°C on a mechanical shaker (200 rpm).

### Construction of PCN-overexpressing and PCN-silenced *P. brasiliensis* yeasts

To obtain a PCN-overexpressing and a PCN-silenced *P. brasiliensis* strains, we used the antisense RNA (AsRNA) strategy and *Agrobacterium tumefaciens* - mediated transformation (ATMT), as described previously by Fernandes *et al*. [7] and Gonçales *et al*. [8]. Briefly, to obtain *P. brasiliensis* strains with the silenced PCN gene, two regions (AS1 and AS2) of the second exon of the PCN gene were amplified and individually inserted into the pCR35-RHO2 vector. The pCR35-RHO2 vectors carrying AS1 or AS2 were used for amplification of the AsRNA cassettes and were individually inserted into the transfer DNA (T-DNA) of the parental binary vector pUR5750. These vectors were used to transform *A. tumefaciens* LBA1100. The transformants were selected in LB medium containing 100 mg/ml of kanamycin and were subsequently used for the transformation of *P. brasiliensis* by co-cultivation with yeasts at a ratio of 1:10 in sterile Hybond N filters (GE Healthcare Life Science, Pittsburgh, PA) in IM solid medium (induction medium) at 28°C for 3 days. After co-culture, the membranes were transferred to liquid BHI medium containing cefotaxime (200 mg/ml). Cells in suspension were incubated for 48 h at 200 rpm at 37°C before being transferred to selective BHI medium containing hygromycin B (100 mg/ml) and cefotaxime (100 mg/ml) at 37°C for 15 days. To obtain the lines with overexpression, the methodology was the same, however, the complete gene was inserted in the vector in the sense. The determination of silencing and overexpression of the clones was verified by qRT-PCR. All mutants were mitotically stable.

### Antifungal drugs

Amphotericin B (AmB), itraconazole (ITZ) and micafungin (MFG) were obtained from the manufacturer (Sigma-Aldrich, MO, USA) and prepared according to the document M27-S4, proposed by Clinical and Laboratory Standards Institute in 2012 with adaptations for *P. brasiliensis*.

### Cloning, expression in *Pichia pastoris*, and purification of recombinant paracoccin

Recombinant PCN was obtained as described by Freitas *et al*. [6]. Briefly, the PCN ORF, cloned into pUC57 vector by Alegre *et al*. [3], was amplified with forward primer 50-CTCGAGATGGCGTTTGAAACCAGATTG-30 and reverse primer 50-GCGGCCGCCCAGCTGCTGGTGCTAAAGC-30. The purified PCR product was cloned into the pGEM-T vector (Promega, Fitchburg, WI, USA), and later into the pGAPzαA vector (Invitrogen, Carlsbad, CA, USA). The pGAPzαA-PCN vector was used for the transformation of the *Pichia pastoris* GS115 strain, as described by Maleki *et al*. [9]. Transformed lines, obtained in YPD selective medium containing Zeocin (100 mg/mL), were confirmed by PCR. The selected clone was cultivated in 300 mL of liquid YPD at 30°C, 220 rpm, for 72 h, for expression and secretion of recombinant PCN. The culture supernatant was collected, dialyzed against PBS (pH 7.2) and concentrated 10 times using centrifugal filtration devices with a 10,000-molecular weight cut-off (Millipore, Darmstadt, Germany). For PCN purification, the culture supernatant was applied in chitin column, manufactured as described by Dos Reis Almeida *et al*. [2]. The product obtained from the chromatography was analyzed on SDS-PAGE and submitted to quality control of biological activity by evaluating its lectin and enzymatic activities.

### Minimum inhibitory concentration (MIC)

Susceptibility tests of *P. brasiliensis* (wt-PCN, ov-PCN, kd-PCN and EV) to the AmB, ITZ and MFG were performed according to De Paula e Silva *et al*. [10] with adaptations. Inoculi were prepared in Roswell Park Memorial Institute (RPMI-1640) medium (Sigma-Aldrich) with l-glutamine, without sodium bicarbonate, supplemented with 2% glucose, and buffered to a pH of 7.0 using 0.165 M morpholine propane sulfonic acid (MOPS; Sigma-Aldrich), to achieve a final concentration in microdilution plates of 0.5 × 10^5^ colony-forming units (CFU)/ml. As quality control, *Candida parapsilosis* ATCC 22019 was tested in the concentration of 2.5 × 10^3^ colony-forming units (CFU)/ml. Work solutions of AmB, ITZ and MFG were tested in concentrations ranged from 0.001955 to 8 μg.ml^−1^, 0.00001525 to 8 μg.ml^−1^, and 0.0156 to 128 μg.ml^−1^, respectively, after adding the inoculum. The plates were incubated at 37°C under agitation of 150 rpm for up to 72 h. The readings were performed using Alamar Blue® (Sigma-Aldrich). The lowest antifungal agent concentration that substantially inhibited the growth of the organism was visually determined at the point at which there was no change in the original blue color of the reagent. Three independent experiments were conducted.

### Minimum fungicide concentration (MFC)

A qualitative analysis of fungal viability was performed, by transferring a portion of the wells to a plate with BHI agar (Kasvi) and incubated at 37°C for 7-10 days. The minimum fungicide concentration was determined as the lowest concentration of the compound that did not allow the growth of any fungal colony on the solid medium after the incubation period. A visual reading was performed [11]. Three independent experiments were performed.

### Analysis of the impact of antifungals on yeast morphology

The impact of the antifungal action on PCN-overexpressing and a PCN-silenced *P. brasiliensis* yeast cells was evaluated by microscopy using an inverted microscope equipped for observation in a bright field (Leica DMI 6000B, Wetzlar, Germany). Yeasts that were not subjected to the action of drugs were used as control.

### Cell wall stress susceptibility

An ideal way to determine the susceptibility to stressors is to inoculate a series of yeast concentrations in the form of spots on plates containing the stressor [12]. For the stress susceptibility analysis, 72-h old *P. brasiliensis* yeast cells (wt-PCN, ov-PCN, kd-PCN and EV) were adjusted to concentration 10^7^, 10^6^ and 10^5^ cells/mL and spotted onto plates with BHI agar (Kasvi) supplemented with different stressor agents at different concentrations as follows: calcofluor (CFW, 20 μg.ml^−1^ and 40 μg.ml^−1^), congo red (CR, 25 μg.ml^−1^ and 50 μg.ml^−1^), sodium chloride (NaCl, 100 mM and 200 mM), hydrogen peroxide (H_2_O_2_, 5 mM and 7,5 mM) and sodium dodecyl sulfate (SDS, 0,005% and 0,010%). Control plates did not contain any stressors. The plates were incubated for 7-10 d at 37°C before being photographed. Additionally, the susceptibility of *P. brasiliensis* strains against cellular stress inducing agents was tested *in vitro* by MIC determination according to the document M27-S4 [13]. Work solutions of CFW, CR, NaCl, H_2_O_2_ and SDS were tested in concentrations ranged from 0,078 to 40 μg.ml^−1^, 0,097 a 50 μg.ml^−1^, 0,781 a 400 mM, 0,097 a 50 mM and 0,0000195 a 0,010%, respectively, after adding the inoculum. The impact of stressors on the yeast cell wall was evaluated by using an inverted microscope equipped for bright field observation (Leica DMI 6000B, Wetzlar, Germany). Yeasts that were not exposed to the action of stressors were used as control.

### Insects

*Galleria mellonela* larvae were kept in a glass container at 25°C in the dark and fed with specific manipulated feed. For the assays, larvae without color changes and with adequate weight (150-200 mg) were selected and kept without food in Petri dishes, at 37°C in the dark, for a period of 24 h prior to infection.

### *Galleria mellonella* infection

Infection of larvae was performed as described by De Lacorte Singulani *et al*. [14]. Prior to injection, an area of the pro-leg was sanitized with 70% alcohol. Then, the larvae were inoculated using a Hamilton syringe (Hamilton Co., Reno, NV), in the last left proleg. Each group of larvae was inoculated with 10 μL of wild-type (wt-PCN), transformants (ov-PCN and kd-PCN) and empty vector (EV) strains of *P. brasiliensis* (5×10^6^ cells/larva). In all assays, a group of uninfected larvae and a group of larvae inoculated with Phosphate-Saline buffer (PBS) were used as controls. For each condition, 8 larvae were used, and each experiment was reproduced three times.

### Survival assay

Infected larvae were incubated in Petri dishes at 37°C and evaluated for 12 days for lack of physical movement (motility) and melanization.

### Fungal burden

At 48 h post-infection, larvae from each group were superficially disinfected with 70% ethanol. Then, through puncture in the abdomen of the larvae, hemolymph samples were collected and diluted in ice-cold PBS with 20 mg/L ampicillin (1:10). Then, 100 μL aliquots of the hemocyte suspension were plated on BHI agar supplemented with 1% glucose and 4% fetal bovine serum (FBS) containing 100 μg/ml Ampicillin. The plates were incubated at 37°C for 10 days. At the end of this period, the colony forming units (CFU) were counted.

### Hemocytes density

At 48 h post-infection with wild-type strains and transformants of *P. brasiliensis*, hemolymph samples were collected by puncturing the larval abdomen and were diluted in ice-cold PBS (1:20). Then, 10 μL aliquots of the hemocyte suspension were counted in a Neubauer hemocytometer.

### *In vivo* resistance assays

At 1 h after infection with *P. brasiliensis*, larvae were inoculated with AmB 0.5 mg/kg in the last right pro-leg. AmB stock solutions were prepared in Dimethylsulfoxide (DMSO) (Labsynth, Diadema, SP, Brazil) and diluted in PBS to a DMSO concentration of 5% as described previously [14]. A group of uninfected larvae was treated with the antifungal alone to assess its toxicity. Larvae were incubated at 37°C and assessed for 7 days for lack of physical movement. Additionally, within 48 h after treatment with AmB, hemolymph samples were collected through puncture in the larvae’s abdomen and hemocytes density was established by microscopy in a Neubauer hemocytometer.

### Histopathological evaluation

After infection (48 h post-infection) and after treatment with AmB (48 h postreatment), the larvae of each group were fixed by immersion in 4% buffered formalin for 24 h. Then, the larvae were preserved in 70% ethanol and longitudinal incisions in the dorsal part were made with the aid of a scalpel. The samples were dehydrated with increasing concentrations of ethanol, washed with xylene, embedded in paraffin, sectioned serially at a thickness of 5 μm and stained with periodic acid Schiff (PAS) (Sigma-Aldrich). Images were acquired on an Olympus microscope comprising the Virtual Slide VS120 system (Olympus, Tokyo, Japan) on a 20× objective and analyzed with FIJI software (ImageJ; NIH, Bethesda, MD, USA).

### Crude extract preparation, RNA Isolation, cDNA Synthesis and qPCR Assays

Cultures of wild and transformant *P. brasiliensis* strains were maintained weekly by subcultures on brain-heart infusion agar (BHI - Duchefa, Netherlands) supplemented with 1% glucose and 4% FBS (Life Technologies) or YPD agar (Yeast extract - Peptone - Dextrose) as described by Pitangui *et al*. [15]. Then, yeast colonies were transferred to YPD broth, incubated at 37° under agitation (150 rpm) for 72 h and collected by centrifugation for 10 min (2300 × g, 4°C). Yeast cells were washed with PBS and total RNA was isolated using Trizol (Life Technologies, Carlsbad, CA, USA) according to the manufacturer’s instructions. The RNA samples quality and concentration were determined by electrophoresis in a 1.2% (w/v) agarose gel with observation of the image in a gel documentation system (Chemidoc MP Imager, Bio-Rad Laboratories, Richmond, CA, USA) and spectrophotometric analysis in NanoDrop® (Thermo Fisher Scientific, Wilmington, DE, USA). The RNA samples were treated with DNAse I (Fermentas, Waltham, Massachusetts, USA) to remove the genomic DNA and, subsequently, the cDNA was synthesized using the cDNA synthesis kit (iScript cDNA Synthesis Kit, Bio-Rad) according to the manufacturer’s instructions. PCR was developed using EVA Green (Bio-Rad) on a CFX96 Real-Time Detection System (Bio-Rad) in reaction volumes of 15 μl under the following conditions: 95°C for 30 s, then 40 cycles of 95°C for 5 s and 60°C for 5 s. Quantification of gene expression was performed by the ΔΔCt method in relation to endogenous controls of α-tubulin and L34. The specific gene primers used for quantitative RT-PCR of fungal cell wall synthesis and degradation markers were: *PbFKS1, PbCHS3, PbCSR1, PbNAG1, PbBGN1, PbBGN2* and *PbAGN*. The sequences of the primers used in the RT-PCR assay are shown in Table 1. The analysis was conducted as previously described [16].

**Table 1.**
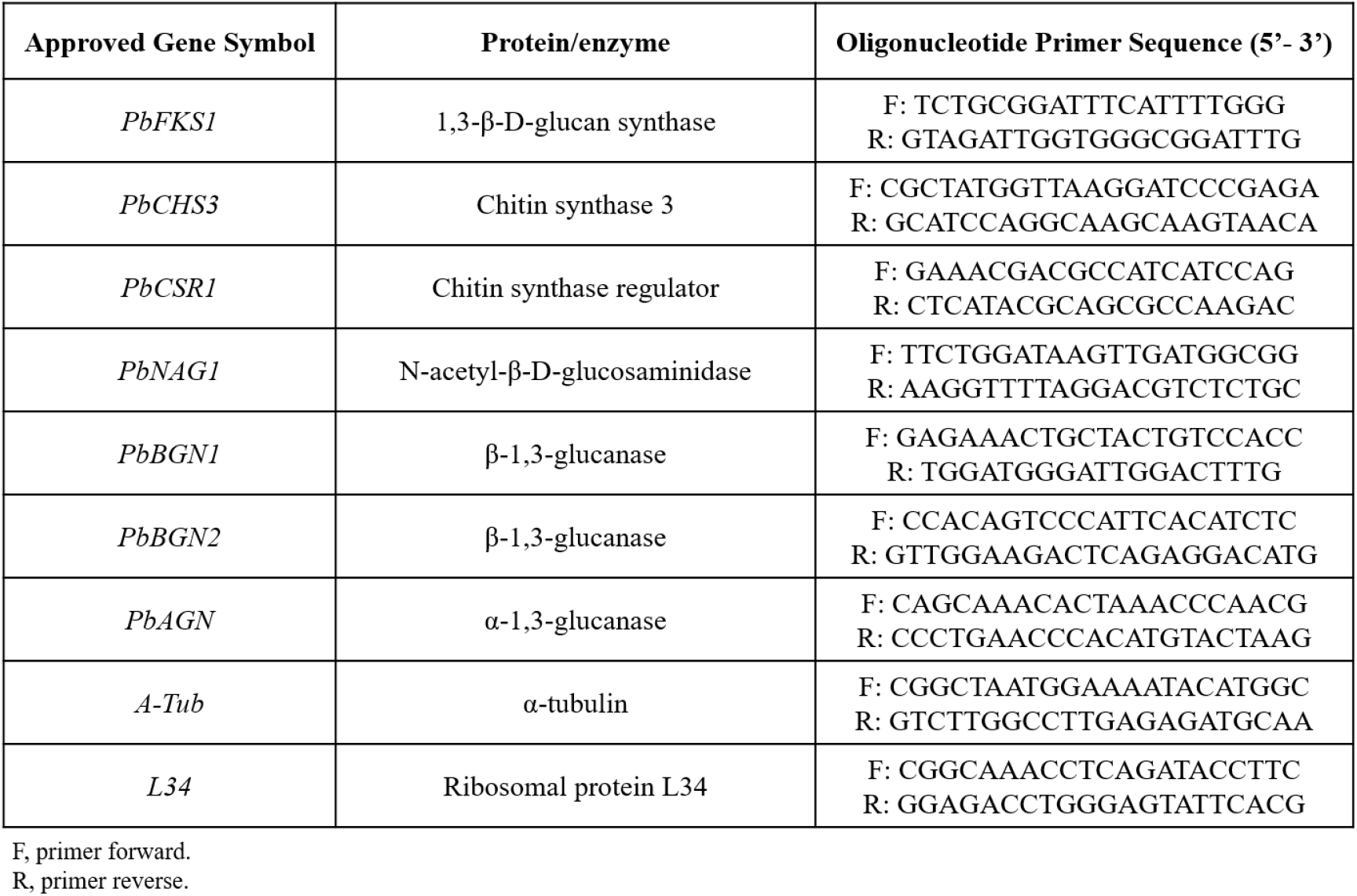
Primers used in this study, and their target genes.

### Statistical Analysis

Data were analyzed using GraphPad Prism v7.00 (GraphPad Software, Inc., La Jolla, CA, USA). Results were expressed as mean ± SD (standard deviation), with observed differences being considered significant when *p* < 0.05 (*). All data are representative of three independent experiments. For the analysis of quantitative RT-PCR data, the One-Way ANOVA test for analysis of variance was performed, with Bonferroni’s multiple comparison test.

## RESULTS

### 1. PCN overexpression correlates with high susceptibility of *P. brasiliensis* to antifungal drugs

To investigate whether PCN could influence the susceptibility of *P. brasiliensis* yeasts to antifungal drugs, we evaluated AmB, ITZ, and MFG lowest concentrations required to inhibit the in vitro growth of yeasts differing in PCN expression (wt-PCN, ov-PCN, and kd-PCN). To ascertain the Minimum Inhibitory Concentration (MIC) parameter, we cultured yeasts from each strain, at equal concentrations (0.5 × 10^5^ colony-forming units (CFU)/mL), in liquid medium. The test drugs, at several concentrations, were added to the cultures. After 72 h, yeasts were stained with Alamar Blue, which allowed to visualize the lowest concentration of each drug required to inhibit the yeasts growth. As shown in Table 2, for each assayed drug, equal or close MIC values were found for wt-PCN and kd-PCN. These values were at least 2 to 4-fold higher than the MIC achieved by ov-PCN yeasts, a disparity indicating that PCN overexpression correlates with a yeast in vitro profile of high susceptibility to all assayed antifungal drugs. Yeasts from wt-PCN and kd-PCN strains displayed low vulnerability to drugs, as shown by the high MIC values determined for all tested drugs.

**Table 2.** Minimum inhibitory concentration (MIC) and minimum fungicidal concentration (CFM) of antifungal drugs against *P. brasiliensis* yeasts from several strains.

|  | AmB |  | ITZ |  | MFG |  |
| --- | --- | --- | --- | --- | --- | --- |
| | MIC ($\mu\text{g.mL}^{-1}$) | MFC ($\mu\text{g.mL}^{-1}$) | MIC ($\mu\text{g.mL}^{-1}$) | MFC ($\mu\text{g.mL}^{-1}$) | MIC ($\mu\text{g.mL}^{-1}$) | MFC ($\mu\text{g.mL}^{-1}$) |
| **Strains** |  |  |  |  |  |  |
| wt-PCN | 0.125 | 0.125 | 0.000488 | 0.000488 | >128 | >128 |
| ov-PCN | 0.0313 | 0.0313 | 0.000122 | 0.000122 | 64 | 64 |
| kd-PCN | 0.125 | 0.25 | 0.000488 | 0.000976 | >128 | >128 |
| EV | 0.125 | 0.125 | 0.000488 | 0.000488 | >128 | >128 |
| AmB: amphotericin B; ITZ: itraconazole; MFG: micafungin; MIC: minimum inhibitory concentration; MFC: minimum fungicidal concentration<br>wt-PCN: *P. brasiliensis* wild-type strain; ov-PCN: PCN-overregulated strain; kd-PCN: PCN-downregulated strain and; EV: empty vector. |  |  |  |  |  |  |

The second parameter used to analyze the yeasts’ susceptibility to the tested drugs was the determination of the Minimum Fungicide Concentration (MFC), which discriminate whether the drug effect verified by MIC has a fungistatic component, besides a fungicidal mechanism. MFC determination was done by transferring part of the wells content of the microplate prepared for MIC determination to a plate with BHI agar. After one week cultured, the MFC was visually determined as the lowest concentration of the drug that completely avoided the growth of any fungal colony on the plate. The CFM value for each antifungal drug determined for yeasts of a certain strain was compared with the respective MIC value. The antifungals AmB, ITZ, and MFG provided CFM equivalent to the MIC values for wt-PCN and ov-PCN yeasts. Concerning the kd-PCN yeasts, the MFC values for AmB and ITZ were 2-fold higher than the respective MIC values. A precise determination of MIC and MFC for MFG was prejudiced by the fact that the highest drug concentration assayed (128 μg.mL^-1^) was not sufficient to completely inhibit the wt-PCN and kd-PCN yeasts growth. Because 64 ug/mL MFG was enough to block the growth of ov-PCN yeasts, we could determine MIC and MFC for this strain, whose values are coincident, and verify that they are at least 2-fold lower than values of MIC and MFC (>128 μg.mL^-1^) attributed for the other yeasts strains.

Our results suggest that a fungistatic component of the tested drugs AmB and ITZ contributed for their apparent better performance when MIC parameter was determined. We suppose that this contribution is little since only a slight difference between the MIC and MFC values was detected for AmB and ITZ.

### 2. PCN-overexpression enhances the *P. brasiliensis* cell wall damage caused by micafungin

We performed yeasts microscopic analyses to evaluate whether antifungal agents impact differently the growth pattern of ov-PCN and kd-PCN yeasts compared to the wt-PCN yeasts. The treatment with AmB or ITZ did not define significant differences in the transformed yeasts compared to the wt-PCN yeasts (data not shown). The yeasts treatment with MFG, in its turn, determined distinct growth patterns on ov-PCN compared to wt-PCN yeasts (Figure 1). Following treatment with 64 μg.mL^-1^ and 128 μg.mL^-1^ MFG, the wt-PCN, and the kd-PCN yeasts have grown by forming aggregates. Otherwise ov-PCN yeasts grew individually, without clustering, a consequence of a more severe damage of the cell wall.

**Figure 1.**
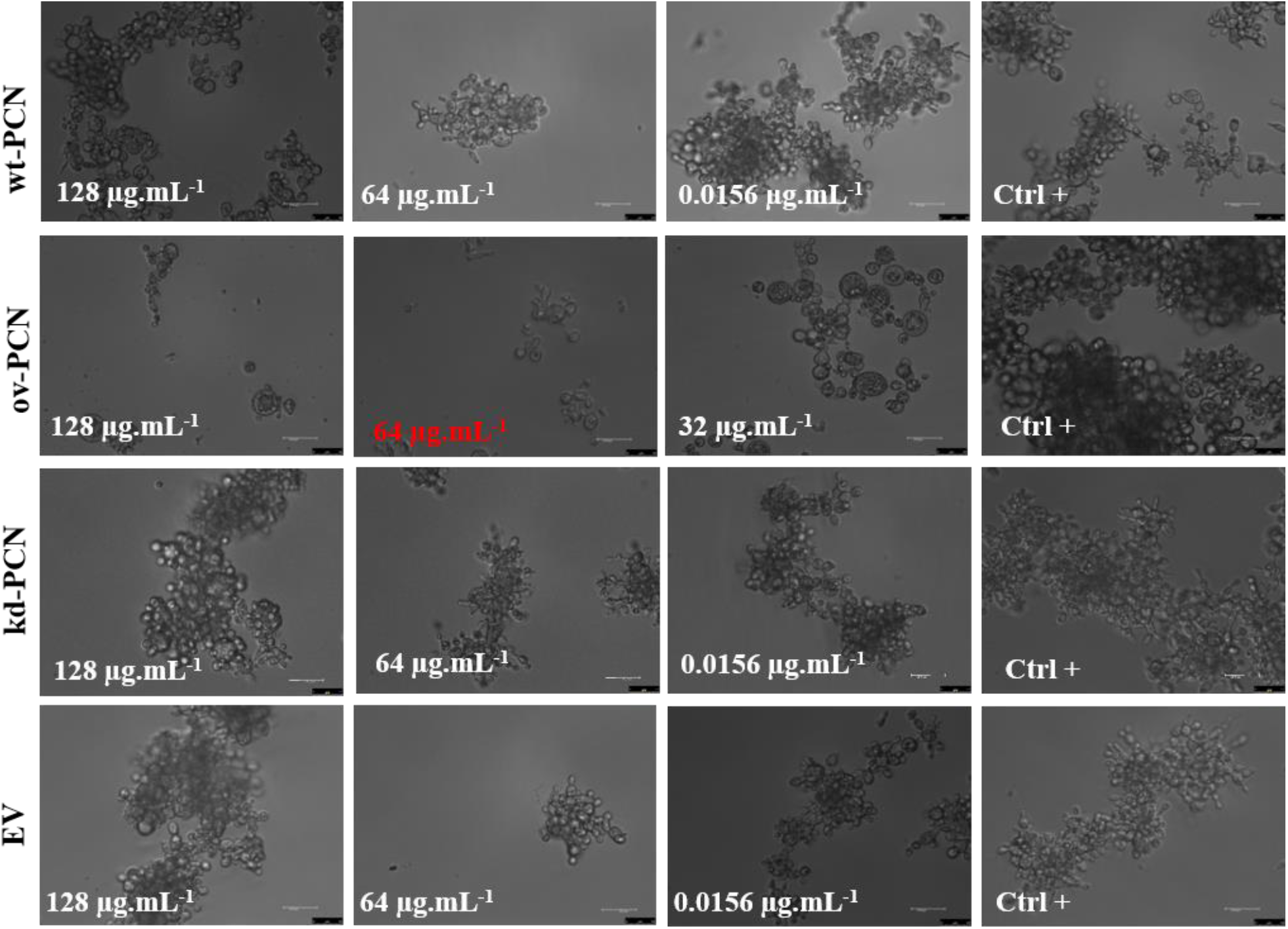
Micafungin influences differently the growth pattern of yeasts from various *P. brasiliensis* strains. Concentration highlighted in red represents the Minimum Inhibitory Concentration (MIC). wt-PCN: *P. brasiliensis* wild-type strain; ov-PCN: PCN-overregulated strain; kd-PCN: PCN-downregulated strain; and EV: empty vector. Ctrl+: Growth control, yeast free from drug effects. 63x objective. Bars, 50 μm.

In addition, the cell wall changes of the PCN-overregulated strain were examined by microscopy after incubation of strain with 64 μg.mL^-1^ MFG. ov-PCN markedly alters the structural integrity of its cell wall after contact with antifungal for 72 h (Figures 2C and D), while wt-PCN, kd-PCN and EV do not exhibit structural damage to the cell wall after incubation with MFG (Figures 2A-B, E-F and G-H).

**Figure 2.**
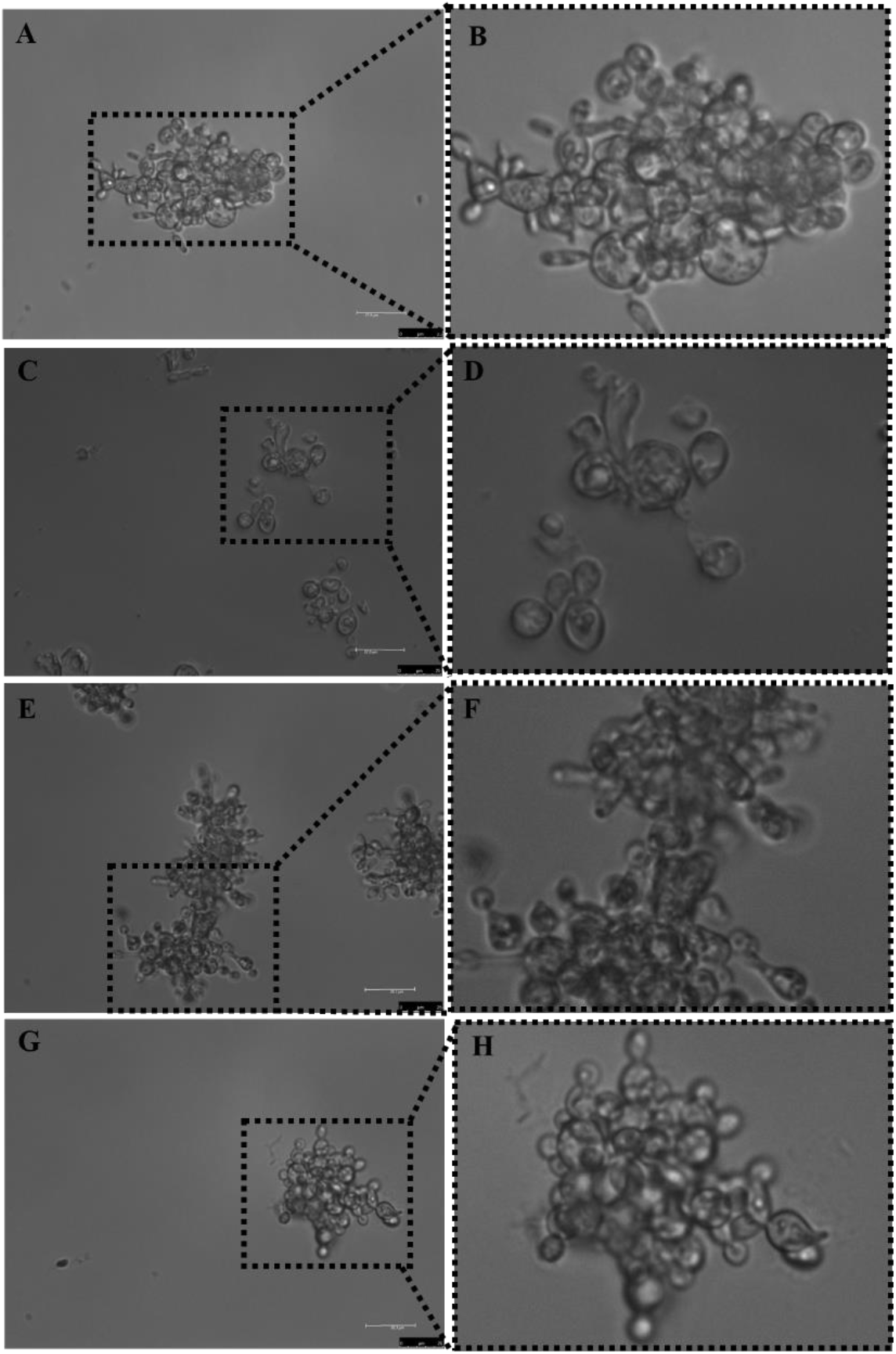
ov-PCN showed disordered cell walls and altered general pattern of yeast organization by the action of micafungin. *P. brasiliensis* yeasts were incubated with 64 μg.mL^-1^ micafungin for 72 h and images were obtained from wt-PCN (*P. brasiliensis* wild-type strain, **A** and **B**), ov-PCN (PCN-overregulated strain, **C** and **D**), kd-PCN (PCN-downregulated strain, **E** and **F**), and EV (empty vector, **G** and **H**). 63x objective and bars, 50 μm in A, C, E and G.

### 3. PCN overexpression enhances the *P. brasiliensis* susceptibility to stress at the cell wall

We evaluated the susceptibility of *P. brasiliensis* yeasts (10^5^, 10^6^, and 10^7^ cells/mL) to agents that induce cellular stress (oxidative, osmotic, and structural stress at the membrane or cell wall level). The following stress agents were employed: Calcofluor (a chitin ligand), Congo Red (a β-1,3-glucan ligand), NaCl (induces osmotic stress, alters ionic homeostasis), H_2_O_2_ (enhances cellular oxidative processes) and SDS (solubilizes cell membrane lipids). Susceptibility was qualitatively estimated by spotting different yeasts′ concentration onto plates with solid medium supplemented with the stressor (at two different concentrations). Yeasts growth was estimated by the visual examination of the fungal colony size on the culture solid medium.

Treatment with 100 mM NaCl did not affect the susceptibility of ov-PCN, kd-PCN and wt-PCN yeasts since the obtained growth profile for each strain was like the given by the respective yeasts’ positive controls (not undergone to cellular stress) (Figure 3). When 200 mM NaCl was used, there was an augmented susceptibility of yeasts 10^5^ concentrated of all strains, except for the kd-yeasts, whose susceptibility was like the verified in the absence of stress. It indicates that when silenced for PCN, the yeasts become more resistant to osmotic stress.

**Figure 3.**
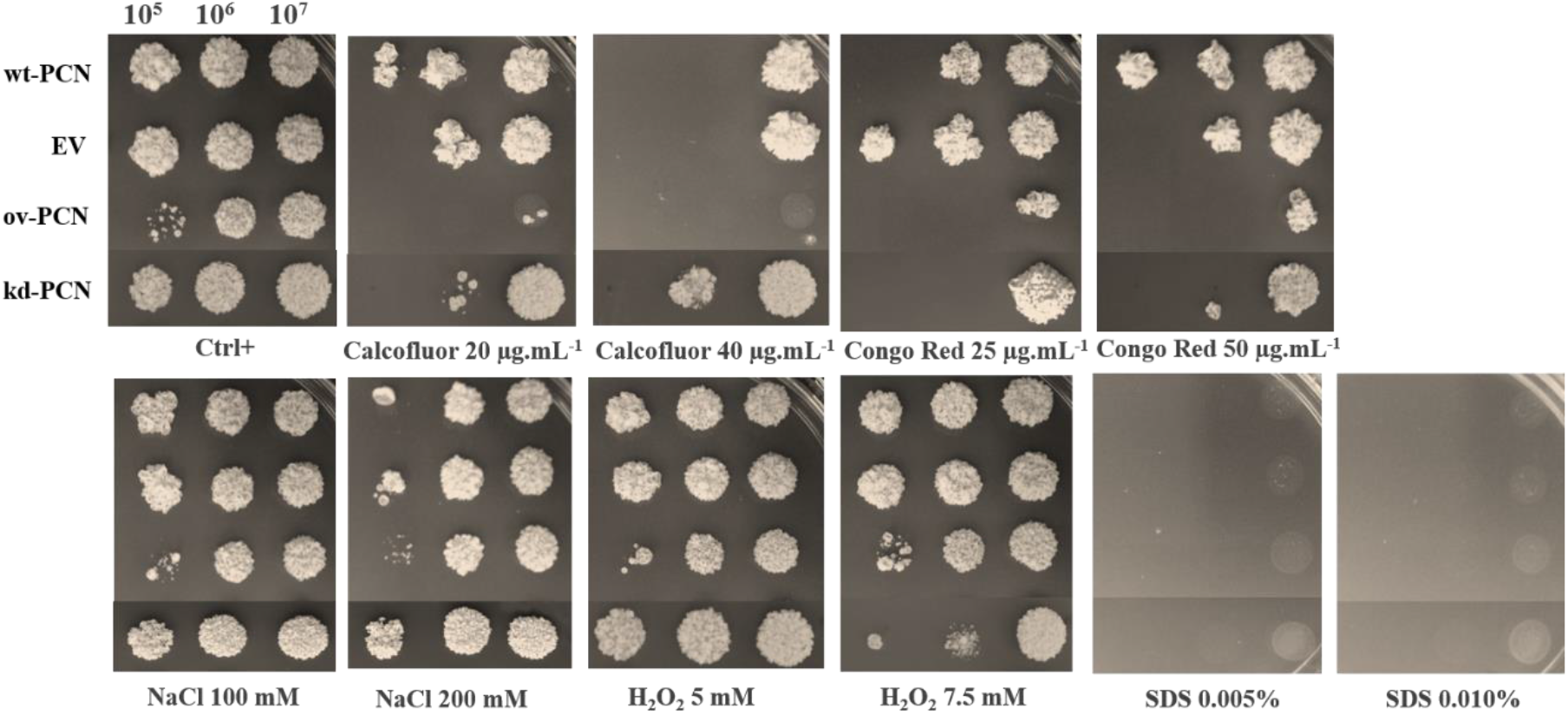
Susceptibility of yeasts from various *P. brasiliensis* strains to cellular stress-inducing agents. The following *P. brasiliensis* strains were assayed: wt-PCN yeasts; ov-PCN yeasts; kd-PCN yeasts; and EV: empty vector. Ctrl+: control of yeasts grown on BHI agar containing 100 μg/ml ampicillin. The plates were incubated at 37°C for 7-10 days for colonies formation. The colonies were visually examined.

By assaying an agent that induces oxidative stress (H_2_O_2_), we verified that, used at 5.0mM, it did not modify the yeasts growth, which stayed like the visualized in the absence of stressor (Ctrl+), for each fungal strain. Nonetheless kd-PCN yeasts (10^5^ and 10^6^ cells/mL) were more susceptible to the effect of 7.5mM H_2_O_2_.

The agent that affected the integrity of fungal cell membrane (SDS) at both tested concentrations prevented the growth of yeasts from all strains.

Yeasts from all strains showed high susceptibility to agents that induce stress at the cell wall level (Calcofluor and Congo Red). It should be emphasized 20 ug/mL Calcofluor was the most potent stressor for the assayed yeasts. Kd-PCN transformant exhibited particularly increased resistance to the chitin ligand (Calcofluor 20 μg.ml^−1^ and 40 μg.ml^−1^) in relation to the profile exhibited by the wild-type strain (wt-PCN). Comparing the strains susceptibility to both stressors of cell wall, it is noteworthy that, compared to wt-yeasts, the ov-PCN and the kd-PCN yeasts exhibited minimal and maximal resistance, respectively, to the assayed stressors.

This observation made clear the yeast susceptibility provided by the PCN overexpression and the yeast resistance given by the PCN silencing.

The antifungal activity of the agents inducing cellular stress - Calcofluor, Congo Red, NaCl, H_2_O_2_, and SDS - was quantitatively evaluated against yeasts of *P. brasiliensis* strains by determining MIC, i.e., the minimal concentration of a particular agent required to inhibit yeasts growth (Table 3). We confirmed that ov-PCN yeasts are more susceptible to cellular stress-inducing agents that act at the cell wall level (Calcofluor and Congo Red) relatively to the susceptibility profile displayed by the wt-PCN yeasts. The MIC value of ov-PCN yeasts toward Calcofluor was 5 μg/mL, while the MIC value for the wild-type strain was 10 μg.mL^-1^. In relation to Congo Red, the susceptibility of ov-PCN yeasts was even higher compared to the wt-PCN strain, with a MIC value <0.097 μg.mL^-1^ for the transformants ov-PCN and 0.390 μg.mL^-1^ for the wt-PCN strain. Considering the other cellular stress-inducing agents (NaCl, H_2_O_2_ and SDS), it is noted that wt- and ov-PCN yeasts exhibit a similar susceptibility profile.

**Table 3.** Minimum inhibitory concentration (MIC) of cellular stress-inducing agents to inhibit growth of *P. brasiliensis* yeasts from several strains.

|  | Calcofluor | Congo Red | NaCl | H<sub>2</sub>O<sub>2</sub> | SDS |
| --- | --- | --- | --- | --- | --- |
| | MIC ($\mu\text{g.mL}^{-1}$) | MIC ($\mu\text{g.mL}^{-1}$) | MIC (mM) | MIC (mM) | MIC (%) |
| **Strains** |  |  |  |  |  |
| wt-PCN | 10 | 0.390 | 200 | 0.390 | 0.00125 |
| ov-PCN | 5 | <0.097 | 200 | 0.390 | 0.00125 |
| kd-PCN | 20 | 1.562 | 400 | 3.125 | 0.00125 |
| EV | 10 | 0.781 | 200 | 0.781 | 0.0025 |
| NaCl: sodium chloride; H<sub>2</sub>O<sub>2</sub>: hydrogen peroxide; SDS: sodium dodecyl sulfate; MIC: minimum inhibitory concentration<br>wt-PCN: *P. brasiliensis* wild-type strain; ov-PCN: PCN-overregulated strain; kd-PCN: PCN-downregulated strain and; EV: empty vector. |  |  |  |  |  |

Conversely, kd-PCN yeasts are more resistant to cellular stress-inducing agents that act at the cell wall (Calcofluor and Congo Red), osmotic stress (NaCl) and stress oxidizing agent (H_2_O_2_) (Table 3). This is reflected in higher MIC values for kd-PCN yeasts than those exhibited by wt-PCN and ov-PCN strains. Toward Calcofluor, the MIC of kd-PCN transformants was 20 μg.mL^-1^, contrasting with the values of 10 μg.mL^-1^ and 5 μg.mL^-1^ for wt- and ov-PCN yeasts, respectively. The susceptibility profile to the β-1,3-glucan ligand (Congo Red) is represented by MIC values of 0.390 μg.mL^-1^, <0.097 μg.mL^-1^ and 1.562 μg.mL^-1^ for wt-PCN, ov-PCN, and kd-PCN yeasts, respectively. These results suggest that PCN silencing has an impact on the composition of cell wall polysaccharides, chitin and β-1,3-glucan, and is associated with a dose-dependent increase in yeast resistance to cell wall stressors (Calcofluor and Congo Red).

### 4. PCN-overexpression increases the *P. brasiliensis* virulence as verified in vivo in a *G. mellonella* model of fungal infection

This study has included an in vivo investigation of the PCN expression effect on the virulence of *P. brasiliensis* yeasts, accomplished in a model developed in the invertebrate *G. mellonella*. Larvae infected with different strains of *P. brasiliensis* (wt-PCN, ov-PCN, kd-PCN, and EV) at 5×10^6^ yeasts/larva had their survival examined at day-12 after infection. The larvae survival was 37.5% among the infected with the wt-PCN, 25% infected with ov-PCN, and 62.5% infected with kd-PCN (Figure 4A).

**Figure 4.**
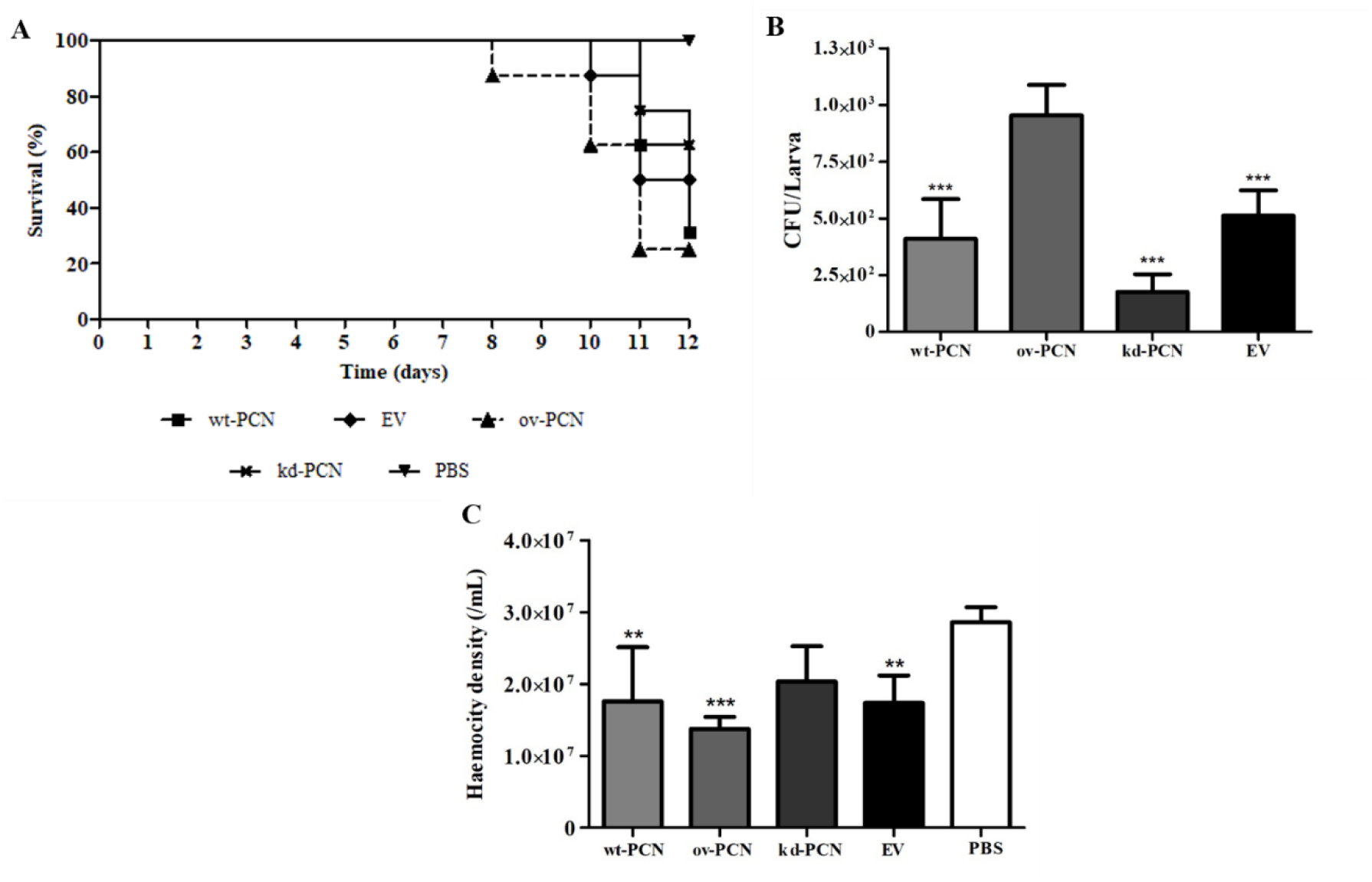
PCN effect on the virulence of *P. brasiliensis* as verified in *G. mellonella* experimental fungal infection. Each *G. mellonella* larvae was infected with 5 × 10^6^ yeasts of wt-PCN, ov-PCN, kd-PCN, or EV (empty vector) (n = 24). Larvae inoculated with PBS were used as negative controls. (**A**) Survival curves of infected *G. mellonella* larvae with *P. brasiliensis* yeasts from different strains. (**B**) CFU recovering 48 h after infection of *G. mellonella* larvae with *P. brasiliensis* yeasts from different strains. *** *p* < 0.0001 *vs*. ov-PCN-infected group. (**C**) Hemocyte density, determined by counting cells in a Neubauer hemocytometer of the hemolymph samples, collected 48 hours after infection with *P. brasiliensis* yeasts from different strains; asterisks indicate statistical significance (*** *p* < 0.0001) relative to the PBS group.

The fungal burden of the *P. brasiliensis* infected larvae evaluated 12-day after infection, through the hemolymph CFU analysis, showed a significantly higher CFU recovery in ov-PCN infected larvae compared to the infected with wt-PCN, kd-PCN, and EV strains. Larvae infected with the three last-mentioned strains exhibited similar fungal loads (Figure 4B).

The third analyzed parameter in the *G. mellonella* model was their hemocyte density, counted in a Neubauer hemocytometer 48 h after infection with *P. brasiliensis* strains. Microscopical examination showed a 2-fold reduction in hemocyte density of larvae infected with wt-PCN, ov-PCN, and EV strains compared to the negative control (PBS). In turn, kd-PCN infected larvae had the hemocyte count like the negative control group (PBS), as shown in Figure 4C.

We investigated whether PCN-silencing and -overexpression influence the anatomopathological aspects of experimental infection by *P. brasiliensis* in *G. mellonella*. Larvae infected with yeasts from one of the studied *P. brasiliensis* strains (wt-PCN, EV, kd-PCN, and ov-PCN) were euthanized 48 h post-infection. Through microscopical examination of body larvae sections, PAS-stained, we verified that in all larvae, independently on the infective *P. brasiliensis* strain, fungal materials were predominantly localized in the periphery of the larvae, close to the cuticle, and the peripheral adipose bodies. We also observed that infection with all *P. brasiliensis* strain determined a similar melanization pattern at the sites of infection. Notably, fungal material intracellularly localized could be detected as cell aggregates of variable sizes, reminding mammal granuloma formation (Figure 5). We found a more significant number of these granuloma-like in larvae infected with ov-PCN yeasts (Figures 5C and 5H), resulting in wide tissue dissemination, with granuloma-like structures occupying a large area of the larval surface. We evidenced this fact by comparison with larvae infected with wt-PCN (Figures 5B and 5G), EV (Figures 5E and 5J), and kd-PCN yeasts (Figures 5D and 5I), in which there are fewer granuloma-like structures, occupying a lesser larval surface.

**Figure 5.**
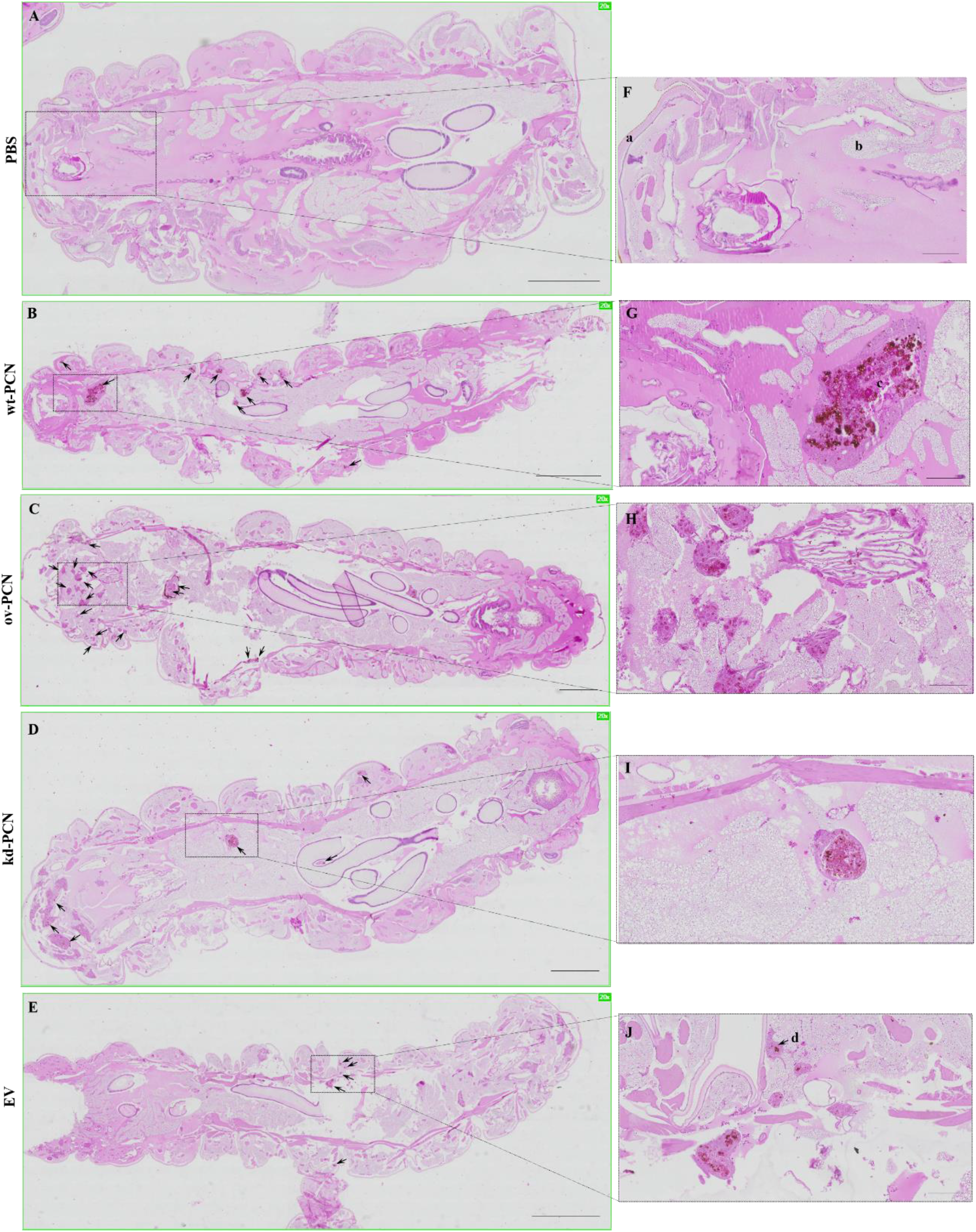
PCN effect on the virulence of *P. brasiliensis* according to histological findings in infected *G. mellonella* larvae. Forty-eight hours after being infected with yeasts from *P. brasiliensis* strains, wt-PCN (B), ov-PCN (C), kd-PCN (D), and EV (E), *G. mellonella* larvae sections were PAS-stained and microscopically examined. Control corresponds to uninfected larvae inoculated with phosphate-buffered saline (PBS) instead of yeasts (A). 20× amplification (A-E). Enlarged images (F-J). Bars indicate 1 mm at A, C, and D; 2 mm in B and E; and 200 μm of F-J. Black arrows indicate *P. brasiliensis* yeast aggregates; a, cuticle; b, adipose bodies; c, granuloma-like structure; d, fungal cells.

### 5. PCN-overexpression increases the susceptibility to Amphotericin B therapy of *G. mellonella* larvae infected with *P. brasiliensis* strains

We assessed the susceptibility profile of *G. mellonella* infected with yeasts from *P. brasiliensis* several strains to the antifungal therapy with Amphotericin B (AmB), the drug of choice for the treatment of severe cases of human paracoccidioidomycosis (PCM).

Figure 6A shows the survival curves of larvae infected with different strains of *P. brasiliensis* (wt-PCN, ov-PCN, kd-PCN and EV) and treated with AmB (0.5 mg/kg), administered 1 h after infection. On day 12 after infection, survival rate of 50% was verified for larvae infected with wt-PCN and EV, 75% for ov-PCN, and 25% for kd-PCN yeasts.

**Figure 6.**
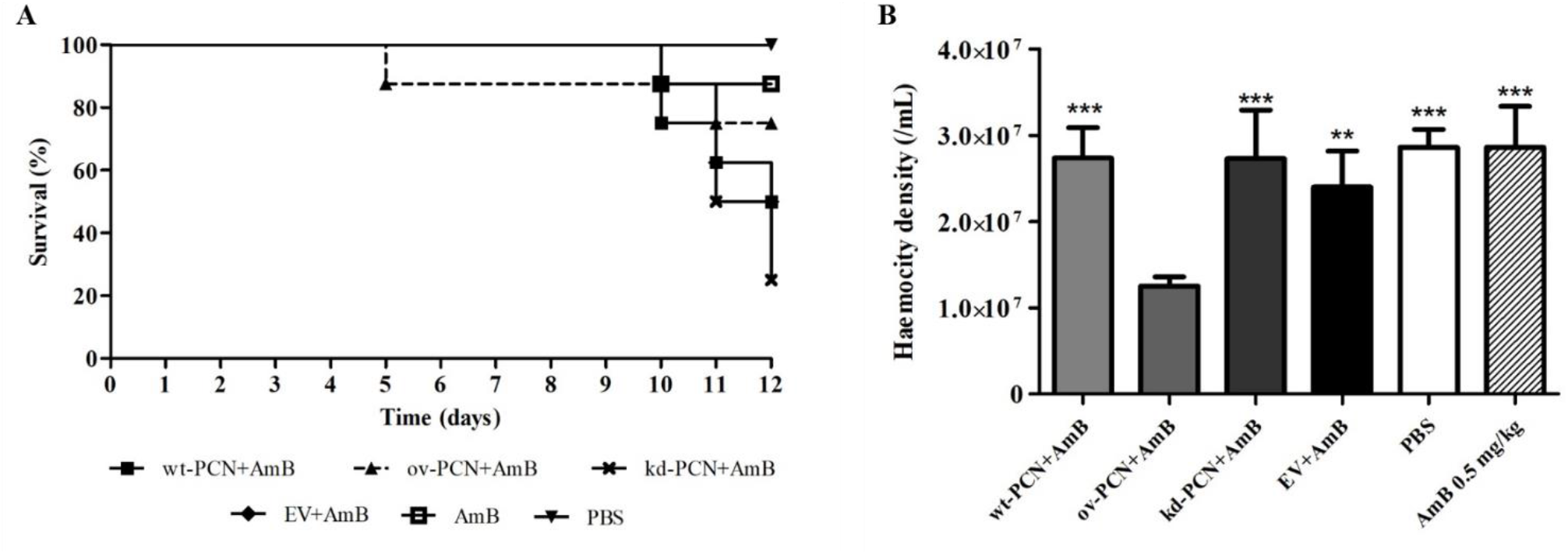
PCN effect on the *P. brasiliensis* susceptibility to Amphotericin B as verified in *G. mellonella* experimental fungal infection. (**A**) Survival curves of *G. mellonella* larvae infected with 5 × 10^6^ yeasts from *P. brasiliensis* different strains and treated at one h post-infection with AmB 0.5 mg/kg. (**B**) Hemocyte density in hemolymph samples collected 48 h after *P. brasiliensis* infection; asterisks indicate statistical significance (*** *p* < 0.0001) *vs*. ov-PCN+AmB. Larvae inoculated with PBS or with AmB were used as controls.

Obeying the same experimental protocol we verified, 48 h after infection, hemocyte densities 2-fold reduced in larvae infected with ov-PCN yeasts. Otherwise, the infection of larvae with wt-PCN, EV, and kd-PCN presented hemocyte density like the exhibited by the controls (PBS and AmB 0.5 mg/kg) (Figure 6B).

We investigated whether *G. mellonella* larvae infected with kd-PCN or ov-PCN *P. brasiliensis* yeasts and treated with AmB present histological evidence of modified susceptibility to the antifungal therapy. Larvae infected with yeasts from one of the studied *P. brasiliensis* strains (wt-PCN, EV, kd-PCN, and ov-PCN) were treated with AmB (0.5 mg/kg) 1 h post-infection and euthanized 48 h post-infection. Figure 7 shows the histological views of *G. mellonella* larvae that were infected with a *P. brasiliensis* strain and treated with AmB.

**Figure 7.**
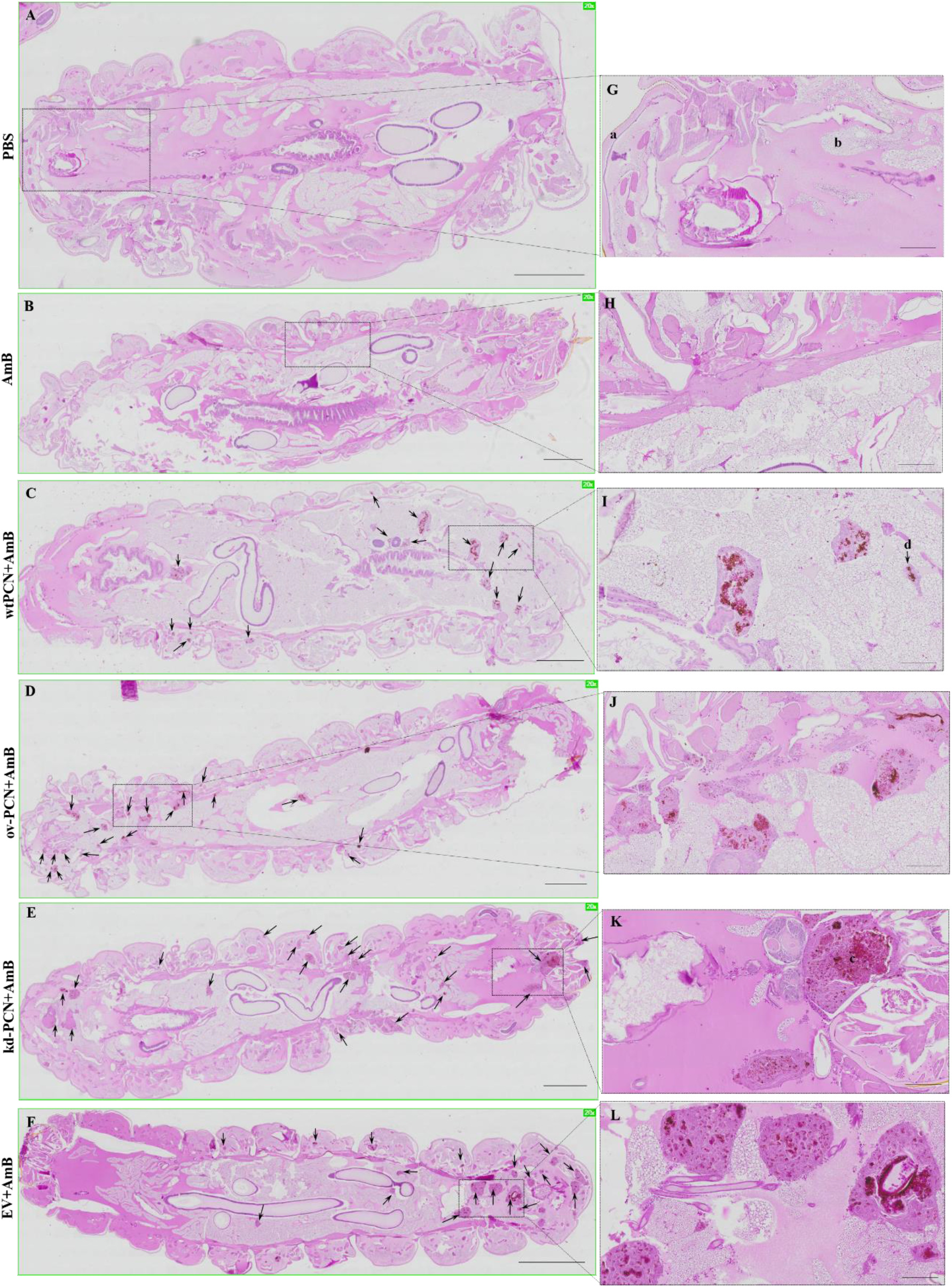
PCN effects on the *P. brasiliensis* yeasts’ susceptibility to Amphotericin B, according to histological findings in infected *G. mellonella* larvae. *G. mellonella* larvae were infected with yeasts from different *P. brasiliensis* strains and treated, 1 hour after, with AmB, 0.5 mg/kg. At 48h post-infection, larvae were euthanized, and body sections were PAS-stained. Images were captured from larvae infected with wt-PCN yeasts(**C**); ov-PCN yeasts (**D**); kd-PCN yeasts (**E**); and EV yeasts (**F**). PBS: uninfected larvae, inoculated with phosphate-buffered saline (PBS) instead of yeasts (**A**). AmB: uninfected larvae inoculated with amphotericin B (0.5 mg/kg) (**B**). 20× amplification (**A-F**). Enlarged views of the images (**G-L**). Bars indicate 1 mm on **A-E**, 2 mm at **F**, and 200 μm at **G-L**. Black arrows indicate *P. brasiliensis* yeast aggregates; **a**, cuticle; **b**, adipose bodies; **c**, granuloma-like structure; **d**, fungal cells.

Larvae infected with kd-PCN yeasts and treated with AmB exhibited a greater number of fungal aggregates and larger extension of tissue injury (Figures 7E and 7K) than the larvae infected with wt-yeasts and treated with AmB (Figures 7C and 7I). The additional comparison with the histological images of larvae infected with kd-PCN yeasts and not treated with AmB (Figures 7D and 7I) showed that the antifungal drug administration was not associated with reduction of host tissue injury. These histological pictures reveal that PCN-silencing augments the yeast resistance to therapy with Amphotericin B.

In their turn, larvae infected with ov-PCN yeasts and treated with AmB exhibited focal aggregates of the fungal material and granuloma-like structures occupying restricted larval areas (Figures 7D and 7J), compared to the larvae infected with wt-PCN yeasts and treated with AmB (Figures 7C and 7I). The additional comparison with larvae infected with ov-PCN yeasts and not-treated with AmB (Figures 7C and 7H) shows that the antifungal drug administration diminishes tissue injury. Together, these results indicate that PCN-overexpression increases the yeast susceptibility to therapy with AmB.

### 6. PCN-overexpression and PCN-knockdown impact differently the relative mRNA expression of markers related to *P. brasiliensis* yeasts cell wall remodeling

Because PCN overexpression aggravates the disease caused by *P. brasiliensis* through a critical effect on fungal cell wall biogenesis, we choose investigating whether PCN influences the relative expression of specific transcripts of markers associated with cell wall remodeling in PCN transformants. We included genes related to cell wall synthesis *PbFKS1, PbCHS3*, and *PbCSR1* since they encode the enzymes 1,3-β-D-glucan synthase, Chs3 (chitin synthase 3), and Csr1 (chitin synthase regulatory protein), respectively. We also evaluated the expression of genes encoding proteins related to cell wall degradation, such as *PbNAG1* (N-acetyl-β-D-glucosaminidase), *PbBGN1* and *PbBGN2* (β-1,3-glucanase), and *PbAGN* (α-1,3-glucanase). After 72h cultured, we found no significant changes in the relative expression levels of the *PbFKS1* and *PbCHS3* genes (related to cell wall synthesis) between the transforming strains (ov-PCN or kd-PCN) and the control (EV) (Figures 8A and 8B). Nonetheless, ov-PCN yeasts showed a diminished relative expression of *PbCSR1* (Figure 8C), a gene encoding one of the putative regulatory proteins of chitin synthase (Csr1). Notably, ov-PCN yeasts, compared to EV control yeasts, hugely reduced the relative expression levels of all genes related to cell wall degradation (*PbNAG1, PbBGN1, PbBGN2*, and *PbAGN*) (Figures 8D, 8E, 8F, and 8G). Compared to kd-PCN, ov-PCN yeasts showed a drastic reduction in the relative expression of *PbNAG1, PbBGN1*, and *PbBGN2* genes (Figures 8D, 8E, and 8F). We normalized data concerning the relative expression levels of the transcripts with the L34 (Figure 8) and A-Tub genes (data not shown); the results were coherent with the data concerning both endogenous controls.

**Figure 8.**
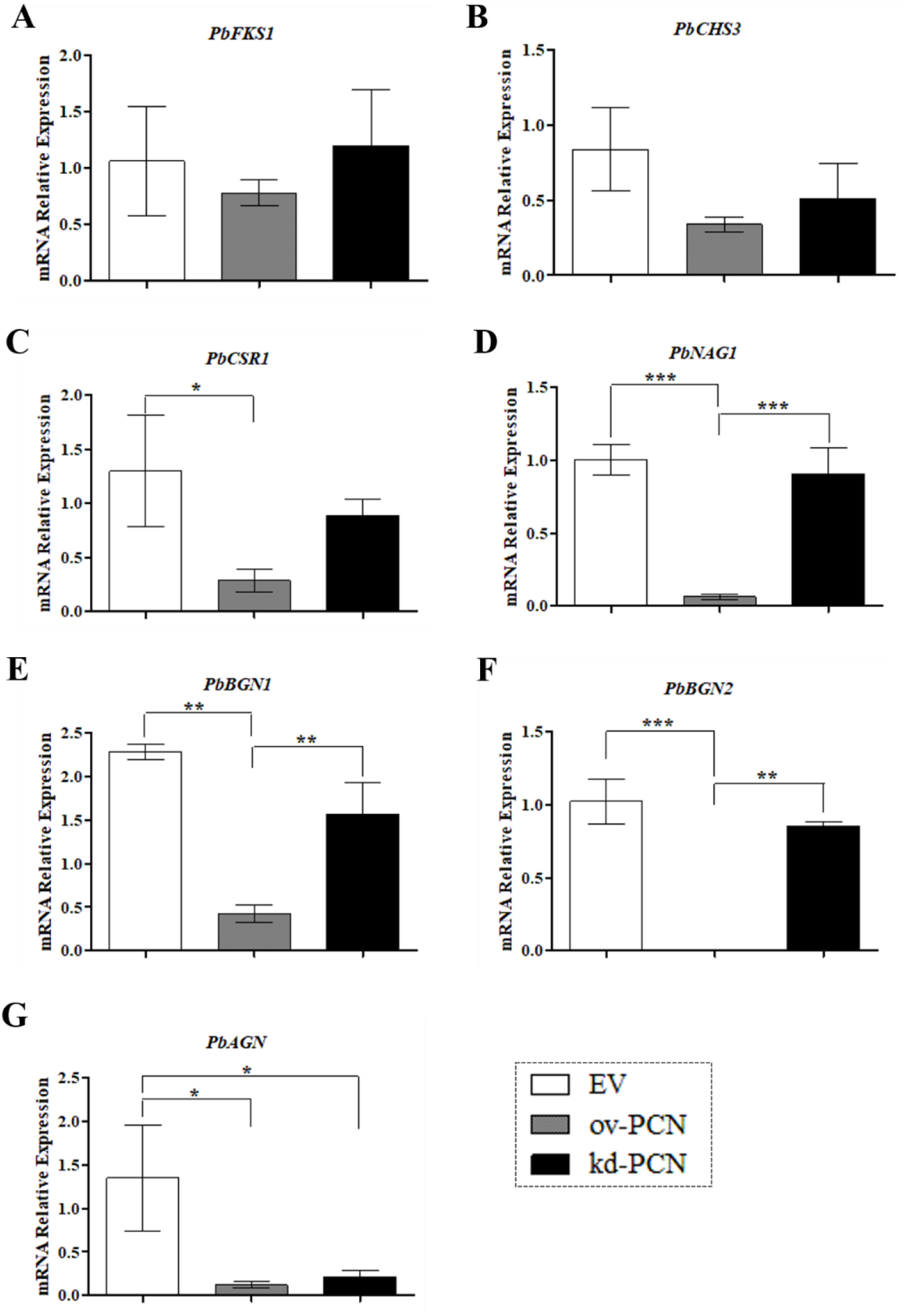
Relative expression of genes related to cell wall remodeling in PCN-transformed *P. brasiliensis* yeasts. The yeasts for RNA extraction were obtained after 72 h cultured. Samples converted to cDNA were analyzed for the relative expression of *PbFKS1* (**A**), *PbCHS3* (**B**), *PbCSR1* (**C**), *PbNAG1* (**D**), *PbBGN1* (**E**), *PbBGN2* (**F**), and *PbAGN* (**G**) by RT-quantitative PCR. The Ct values of the transcripts were normalized to the relative expression of the endogenous control L34. Results are expressed as mean ± SD, and the relative expression levels were compared between yeast transformants (ov-PCN or kd-PCN) and the EV-control group. \**p* < 0.05, \*\**p* < 0.001, and \*\*\**p* < 0.0001.

## DISCUSSION

We demonstrate that PCN-overexpressing *P. brasiliensis* yeasts (ov-PCN) are more susceptible to antifungal agents (AmB, ITZ, and MFG) than wild-type yeasts (wt-PCN). This assertion reflects the lower MIC values for ov-PCN than wt-PCN yeasts, undergone the action of antifungal drugs. The microdilution assay used for MIC determination had its quality controlled by establishing the MIC values for *C. parapsilosis* under the AmB and ITZ action; the obtained values were consistent with the CLSI standardization [13] (Data not shown).

We verified that ov-PCN yeasts are more melanized than wt-PCN yeasts. Previously, authors considered melanization a virulence factor that reduces *P. brasiliensis* susceptibility to AmB and azole drugs [17]. Nonetheless, we saw that *P. brasiliensis* yeasts overexpressing PCN, although verified as more melanized cells, are more susceptible to antifungal drugs.

The fungistatic activity of echinocandins (MFG) results from synthesis blockage of the cell wall component β-1,3-glucan [18]. The content of β-1,3-glucan in the wall of *P. brasiliensis* yeasts is significantly lower (3.9 to 10.6%) than the content in mycelial forms (20.2 to 31.4%) [18], justifying the minimal effect of echinocandins on yeasts. We verified that ov-PCN *P. brasiliensis* yeasts are more susceptible to MFG effects since they exhibit a significantly lower MIC (64 μg.ml^−1^) than the displayed by wt-PCN yeasts (>128 μg.ml^−1^). This observation has generated the question of whether β-1,3-glucan content is higher in the cell wall of ov-PCN than in wt-PCN yeasts.

Herein, we clarified that kd-PCN and wt-PCN yeasts present similar susceptibilities to antifungal agents that act in the fungal membrane or cell wall (AmB, ITZ, and MFG). The technology to silence PCN in *P. brasiliensis* yeasts utilizes antisense RNA (AsRNA) and genetic transformation via *A. tumefaciens* (*A. tumefaciens*-mediated transformation, ATMT) [7]. It generated clones 40-60% reduced in the levels of PCN relative expression (mRNA) compared to wt-PCN yeasts. We hypothesize that, although the reduced levels of PCN expression were sufficient to diminish the separation among yeast cells, making them more clustered [7], they could not confer to kd-PCN yeasts a distinct resistance when exposed to antifungal drugs.

Through light microscopy, we demonstrated that wt-yeasts and PCN transformants did not alter their morphology when exposed to AmB and ITZ. Nonetheless, incubation with 64 μg.ml^-1^ and 128 μg.ml^-1^ MFG affected the ov-PCN yeasts’ growth and morphology and prevented their clustering, compared to the cellular organization pattern shown by the other strains. This observation is consistent with the fact that AmB and ITZ are drugs acting on the fungal cell membrane, while MFG targets the cell wall, inhibits β-1,3-D-glucan synthase, affects the cell wall composition (β-1,3-glucan) and promotes cell lysis. Our data confirmed that ov-PCN yeasts had augmented β-1,3-glucan content in the cell wall compared to *P. brasiliensis* wt-yeasts content (3.9 to 10.6%), as previously reported [19]. The hypothesis herein built assumes that the MFG inhibitory effect on ov-PCN transformed yeasts is due to increased β-1,3-glucan content in their cell wall.

Our investigation of the effect of stress-inducing agents in the cell wall of *P. brasiliensis* transformants showed that ov-PCN yeasts are more susceptible to the action of Calcofluor and Congo Red than the wild-type yeasts, as demonstrated in qualitative and quantitative susceptibility analyses. We showed an augmentation of the content of polysaccharides that make up the cell wall in ov-PCN yeasts. It is supported by the Selvaggini *et al*. [20] demonstration that augmented fungal hypersensitivity to Calcofluor is closely associated with high chitin content in the cell wall. Then we suggest that yeasts that overexpressed PCN, probably through increased chitinolytic activity, displayed cell wall weakening, an alteration that activates compensatory mechanisms to increase chitin and β-1,3-glucan contents. Augmentation of cell wall polysaccharides accounts for the yeasts’ hypersensitivity to Calcofluor and Congo Red. Contrarily, kd-PCN yeasts were more resistant to the Calcofluor action due to the reduced PCN relative expression, which impacts the fungal cell wall organization, biosynthesis, and remodeling. Briefly, the modified content of polysaccharides in the cell wall of kd-PCN yeasts results in resistance to Calcofluor.

In the last decade, *G. mellonella* larvae became a convenient host model for experimental PCM [14, 21-25], which attends to ethical and economic requirements for good animal experimentation practices. Then, we used this larval model to compare host damages caused by inoculation of wild-type yeasts and PCN-transformed yeasts. A comparison of the larvae infections showed that ov-PCN are the most virulent yeasts. They led to a host diminished resistance, as indicated by decreased survival rate, higher tissue fungal load, and reduced hemocyte density, confirming previous results from mice infected with the same *P. brasiliensis* strains [8]. The wt-PCN yeasts were the second most virulent strain. Otherwise, the kd-PCN yeasts virulence in larvae was minimal, differing vastly from the verified in infections with ov-PCN and wt-PCN yeasts [7]. We previously demonstrated a similar virulence pattern for the same fungal strains in mice.

In this case, the murine host minimized the virulence of kd-PCN yeasts. At the same time, mice resistance increased, as revealed by the detection of the highest larval survival rate, reduced fungal load in larval tissue, and increased larval density of hemocytes. We also took advantage of the larval infection model to investigate the interference of the treatment with AmB on the infection with wt-PCN, ov-PCN, and kd-PCN yeasts. A studied parameter was the survival of the infected larvae treated or not with AmB. Maximal sensitivity to the treatment with AmB was given by ov-PCN yeasts since infection with them resulted in the highest larvae survival (75%). In contrast, infections with wt-PCN and EV yeasts treated with AmB provided 50% survival. The kd-PCN yeasts displayed the lowest susceptibility to treatment with AmB since we verified a 25% larval survival.

The behavior displayed by PCN-transformed yeasts when infecting mice and *G. mellonella* is probably associated with PCN activities such as: (i) it affects the separation among yeast cells, and (ii) influences the size of the chitin fragments released during the host-fungus interaction. Fernandes *et al*. [7] demonstrated that PCN-silencing does not affect yeast cells’ viability and growth; it even reduces the separation between yeast cells. Interestingly, kd-PCN yeasts are more clustered than the wt-PCN ones. Probably agglomeration gives protection to the yeasts, favoring resistance to antimicrobial agents. Conversely, ov-PCN yeasts exhibit low adhesion among yeast cells, a fact attributed to augmented PCN chitinase activity, which promotes intense chitin degradation and decreases the thickness of the fungal cell wall, culminating with fragility and augmented susceptibility to antifungal drugs. Gonçales *et al*. [8] emphasized that the overexpression of PCN, thanks to its N-acetylglucosaminidase activity, promotes chitin hydrolysis to tiny fragments, responsible for increasing the macrophages production of IL-10 and M2-type cell polarization, responses characteristically found in severe and disseminated forms of PCM. This hypothesis may explain how ov-PCN yeasts led to *in vivo* exuberant pathogenicity.

We also supposed that the exacerbated virulence of ov-PCN yeasts could potentiate the AmB cytotoxic effect, a circumstance that would reduce the larval hemocyte density. Infection with the ov-PCN yeasts followed by treatment with AmB has maintained 75% larval survival over a 12-days observation period. Data reported by De Lacorte Singulani and co-authors [14] are like ours when correlating survival curve *vs*. hemocyte density in larvae. The larvae infected with Pb18 yeasts (equal to wt-PCN, 5 × 10^6^ cells/larva) and treated with AmB (2 mg/kg) displayed a 88% survival rate. Despite that, the AmB treatment (2 mg/kg) 48 h after the larval infection did not induce recruitment of hemocytes, whose density was like the verified in the control group larvae, which received PBS instead of the antifungal drug.

We investigated the PCN effect on the anatomopathological features of the *G. mellonella* larvae infection with *P. brasiliensis* yeasts. We verified that PCN impacts the virulence of the yeasts because the larval infection with ov-PCN yeasts showed a high number of fungal aggregates and disseminated tissue lesions. In opposition, larval infection with kd-PCN yeasts resulted in fewer yeast aggregates and less extensive tissue injuries. Notably, PCN overexpression further impacted the *P. brasiliensis* yeasts’ susceptibility to the antifungal treatment, as demonstrated by the fewer granuloma-like structures spread in the tissues of larvae infected with ov-PCN yeasts and treated with AmB. A result of such finding is the higher frequency of tissue yeast aggregates and extended tissue damage in larvae infected with kd-PCN yeasts and treated with AmB. Then, studies performed in the *G. mellonella* model of PCM indicate that the PCN expression degree in *P. brasiliensis* yeasts impacts the *in vivo* fungal virulence and its susceptibility to the antifungal drug AmB.

At the transcriptional level, PCN impacts the expression of *P. brasiliensis* genes related to cell wall remodeling, as demonstrated by a reduced *PbCSR1* relative expression in cultured ov-PCN yeasts. *PbCSR1* is a gene that encodes Csr1, one of the three reported putative fungal chitin synthase regulatory proteins (Csr1, Csr2, and Csr3). They are relevant but not essential for yeasts’ viability [26]; Csr1 importance is due to its ability to interact with Chs3. It is done through a prenylation site (CaaX motif), revealed by Csr1 sequence analysis, which indicates its ability to establish protein-protein interaction, probably with Chs3. Grabińska and co-authors reported this interaction as significant for Chs3 activity in *Saccharomyces cerevisiae* [27]. We confirmed the presence of a putative chitin synthase regulator (Csr1) in *P. brasiliensis* yeasts, which has motivated us to propose that it plays a similar role to the exerted by its analog in *S. cerevisiae* [26], which is essential for the Chs3 activity. We have mentioned already that the overexpression of PCN, a protein with chitinolytic activity, results in low thickness and fragility of the cell wall of *P. brasiliensis* yeasts. This modification, in turn, activates compensatory mechanisms capable of increasing the chitin content, evidenced herein by the hypersensitivity of ov-PCN yeasts to calcofluor action. Transcriptionally, we detected that PCN overexpression does not interfere with the relative expression of *PbCHS3* and that the transformant has a low relative expression of *PbCSR1*. Our data are consistent with previous observations in other fungi species, which showed that transcription levels of genes related to cell wall remodeling do not always correlate with the yeast chitin content [28]. The lack of a direct relationship between the measured transcript level and the recovered enzymatic activity occurs because chitin synthase activity regulation occurs at both transcriptional and post-transcriptional levels [29]. Niño-Vega et al. [28] reported that mRNA levels for genes encoding several *P. brasiliensis* chitin synthases (*PbrCHS1, PbrCHS2, PbrCHS4*, and *PbrCHS5*) were higher in hyphae than in yeast. However, yeast cells contain more chitin than mycelial cells. Thus, our data regarding similar levels of *PbCHS3* expression in ov-PCN and wt-yeasts may be explained by genome analysis. It reveals seven chitin synthases in *Paracoccidioides* [30], while we evaluated only the levels of *PbCHS3* relative expression. Notably, ov-PCN yeasts adopted mechanisms to compensate for the “fragility” of the cell wall, manifested by the reduced relative expression of genes associated with cell wall degradation (*PbNAG1, PbBGN1, PbBGN2*, and *PbAGN*). The hydrolytic enzymes, β-1,3-glucanases and chitinases, deserve emphasis because they participate in morphogenetic events to fragilize the cell wall structure. Beyond chitinases, N-acetyl-β-D-glucosaminidase should be highlighted since it is a glycosyl hydrolase that promotes efficient chitin degradation [31]. We show that PCN overexpression results in low production of N-acetyl-β-D-glucosaminidase, β-1,3-glucanases, and α-1,3-glucanase. Our data reinforce the hypothesis that poor expression of a gene related to cell wall degradation in ov-PCN yeasts may provide a mechanism, at the transcriptional level, to compensate weakening of the cell wall, promoting a reduced degradation of polysaccharides that make up the yeast cells wall, i.e., chitin, β-1,3-glucan, and α-1,3-glucan.

Our study brings new insights into the impact of endogenous PCN on the *P. brasiliensis* yeasts’ virulence *vs*. susceptibility to antifungal drugs, the fungal biology, and the relationship of the yeasts-host cells. PCN overexpression increases the virulence of *P. brasiliensis* yeasts, decreases the *in vitro* and *in vivo* yeasts’ resistance to antifungal therapy, and influences the cell wall remodeling by reducing the relative expression of genes accounting for cell wall degradation. Meanwhile, PCN silencing minimized the yeasts’ virulence and augmented fungal resistance to the interaction with cells of an alternative invertebrate host, as established by infecting *G. mellonella* larvae with transformed yeasts relatively to the PCN endogenous expression.

## Acknowledgements

We thank Érica Vendruscolo, Patrícia E. Vendruscolo and Sandra M. O. Thomaz for technical support.

